# Incidental reinstatement of faces and scenes in medial temporal lobe subregions during word recognition

**DOI:** 10.1101/2021.09.14.460299

**Authors:** Heidrun Schultz, Tobias Sommer, Jan Peters

**Author notes:** **Corresponding author:** Heidrun Schultz, Max Planck Institute for Human Cognitive and Brain Sciences, Stephanstr. 1A, 04103 Leipzig, Germany.

## Abstract

During associative retrieval, the brain reinstates neural representations that were present during encoding. The human medial temporal lobe (MTL) with its subregions hippocampus (HC), perirhinal cortex (PRC), and parahippocampal cortex (PHC) plays a central role in neural reinstatement. Previous studies have given compelling evidence for reinstatement in the MTL during explicitly instructed associative retrieval. High-confident recognition may be similarly accompanied by recollection of associated information from the encoding context. It is unclear, however, whether high-confident recognition memory elicits reinstatement in the MTL even in the absence of an explicit instruction to retrieve associated information. Here, we addressed this open question using high-resolution fMRI. Twenty-eight male and female human volunteers engaged in a recognition memory task for words that they had previously encoded together with faces and scenes. Using complementary uni- and multivariate approaches, we show that MTL subregions including the PRC, PHC, and HC differentially reinstate category-specific representations during high-confident word recognition, even though no explicit instruction to retrieve the associated category was given. This constitutes novel evidence that high-confident recognition memory is accompanied by incidental reinstatement of associated category information in MTL subregions, and supports a functional model of the MTL that emphasises content-sensitive representations during both encoding and retrieval.

## 1. Introduction

Recognition memory – the ability to distinguish previously encountered from novel items – critically depends on the medial temporal lobe (MTL), including the hippocampus (HC), perirhinal (PRC), parahippocampal (PHC), and entorhinal cortex (EC) (Eichenbaum et al., 2007). The individual functions that these subregions serve in recognition memory remain a subject of some debate (Wixted and Squire, 2011; Bird, 2017). One model that aims to account both for behavioural observations and their underlying neural substrate is the dual-process signal detection model (DPSD): In this view, two complementary processes contribute to recognition: Familiarity is a signal detection process resulting in graded recognition confidence, supported by PRC, whereas recollection is a threshold process resulting in high recognition confidence, and involves retrieval of associated information from the encoding context, supported by HC and PHC (Eichenbaum et al., 2007; Yonelinas et al., 2010).

Recent work has integrated such process-based views with more content-based accounts of MTL functioning (Davachi, 2006; Eichenbaum et al., 2007). The latter are based on connectivity studies in non-human primates and rodents (Suzuki and Amaral, 1994a, 1994b; Burwell and Amaral, 1998a; Lavenex and Amaral, 2000). Here, the PRC, anatomically connected to the ventral visual stream, processes items, e.g. objects or faces, thereby contributing to familiarity. Meanwhile, the PHC, anatomically connected to the dorsal visual stream, processes spatial context memory, thereby providing the context information underlying recollection. The HC, exchanging information with both streams via anterolateral and posteriormedial subregions of the EC (alEC, pmEC) (Schultz et al., 2015), supports recollection in a content-agnostic manner (Davachi, 2006; Eichenbaum et al., 2007). MTL connectivity in humans is comparable to animals (Zeineh et al., 2012; Maass et al., 2015; Navarro Schröder et al., 2015). Indeed, a number of human functional magnetic resonance imaging (fMRI) studies have demonstrated sensitivity of the PRC to objects, or faces, and of the PHC to spatial or scene information during both perception/encoding (Awipi and Davachi, 2008; Litman et al., 2009; Staresina et al., 2011; Schultz et al., 2021) and associative retrieval (Schultz et al., 2012, 2019; Staresina et al., 2012, 2013; Mack and Preston, 2016) (for an overview, see Robin et al., 2019). Similar content-based dissociations have been demonstrated between alEC and pmEC for faces/objects and scenes/spatial information, respectively (Schultz et al., 2012, 2015; Reagh and Yassa, 2014; Navarro Schröder et al., 2015; Berron et al., 2018).

Importantly, content-specific neural representations during retrieval overlap with representations during the original encoding episode (Danker and Anderson, 2010) (but see Favila et al., 2020). This so-called neural reinstatement of the encoding context is thought to underlie the subjective impression of re-experiencing an episode that accompanies recollection, but not familiarity (Eichenbaum et al., 2007; Danker and Anderson, 2010). Indeed, the degree of reinstatement is associated with objective accuracy (Gordon et al., 2014; Liang and Preston, 2017) as well as subjective vividness of the retrieved memory (Kuhl and Chun, 2014; St-Laurent et al., 2015; Bone et al., 2020), and interrupting early reinstatement through transcranial magnetic stimulation decreases memory performance (Waldhauser et al., 2016).

Content-specificity during memory retrieval has largely been investigated using paradigms that emphasise intentional associative retrieval (Schultz et al., 2012, 2019; Staresina et al., 2012, 2013; Mack and Preston, 2016), e.g. by presenting a cue that was previously paired with an object or scene, and asking participants to retrieve the object or scene from memory (Schultz et al., 2019). Such intentional cued retrieval paradigms are not necessarily comparable to recognition memory. In a recognition paradigm, the task is to judge whether a given item has been previously encountered, and participants are typically asked to qualify their old/new judgments e.g. by rating their confidence (Yonelinas et al., 2010). Importantly, these confidence ratings refer to recognition confidence for the item itself, rather than any associated information that was present during encoding. Since recollection is a threshold-process assumed to selectively lead to high-confidence recognition (Yonelinas et al., 2010), the contributions of recollection and familiarity can be estimated from the asymmetry of the resulting receiver-operating characteristic (ROC) curve (Dunn, 2010; Yonelinas et al., 2010). Recollection is furthermore assumed to involve retrieval of the encoding context, accompanied by neural reinstatement of associated memory content (Eichenbaum et al., 2007; Yonelinas et al., 2010). It follows that items that are recognised with high confidence ought to be accompanied by neural reinstatement of the encoding context, even in the absence of an explicit instruction to retrieve associated information. However, we are not aware of any studies investigating this proposition in subregions of the MTL.

Here, we test this open question, using distinct categories (faces, scenes) to track content representations in the MTL during perception and recognition. On the first day of the study, twenty-eight participants underwent fMRI while viewing a total of 120 faces and scenes (ten exemplars per category, six presentations per exemplar). The participants’ task was to respond to flickers in the image presentation as quickly as possible to win a small reward (scanned perception phase^1^). Next, they learned a list of 260 words, each presented once while paired with one of the faces or scenes, with the task of combining each pair into a single mental image (unscanned encoding phase). The next day, participants returned for a recognition task of the words only, including all 260 target words from the previous day as well as 130 distractor words (scanned recognition phase). For each word, participants rated their confidence that it was old or new. Importantly, there was no instruction to retrieve the associated face or scene. Finally, they solved a source memory task, in which they responded for each word whether it had been paired with a face or a scene the day before (unscanned source phase). For the behavioural analysis, we summarised memory performance for words previously associated with either face or scenes using both model-based (recollection, familiarity) and model-free (corrected recognition, source accuracy) measures. For the fMRI analysis, we tested for i) category sensitivity during face/scene perception, and ii) category reinstatement of the associated faces/scenes during word recognition within participant-specific MTL subregions of interest (ROIs). To this end, we utilised a set of complementary analyses. For the perception phase, we tested for differences in the mean univariate response of each ROI to face and scene perception, and furthermore established multivariate discriminability of faces vs. scenes by training and testing a face/scene classifier on the perception data in a leave-one-run-out fashion. For the recognition phase, we again characterised each region’s univariate response profile to words previously associated with faces or scenes that were recognised with high or low confidence. Critically, we tested for neural reinstatement during word recognition by training a multivariate face/scene classifier on the perception phase, and testing it on words that were previously associated with faces or scenes and recognised with high confidence during the recognition phase. As there was no perceptual overlap between the phases, the classifier performance can only be driven by reinstatement of the face/scene encoding context. As recollection is thought to involve the reactivation of context information and lead to high-confident recognition judgments (Eichenbaum et al., 2007; Yonelinas et al., 2010), we expected words that were recognised with high confidence to be accompanied by neural reinstatement of the encoding category.

## 2. Results

### 2.1 Behavioural results

Overall, analyses of the behavioural data confirmed i) above chance performance in both the recognition and source phases, and ii) critically, no significant differences between words previously associated with faces (wordsF) and scenes (wordsS) (see Figure 1 for overview of the analysed measures).

**Figure 1.**
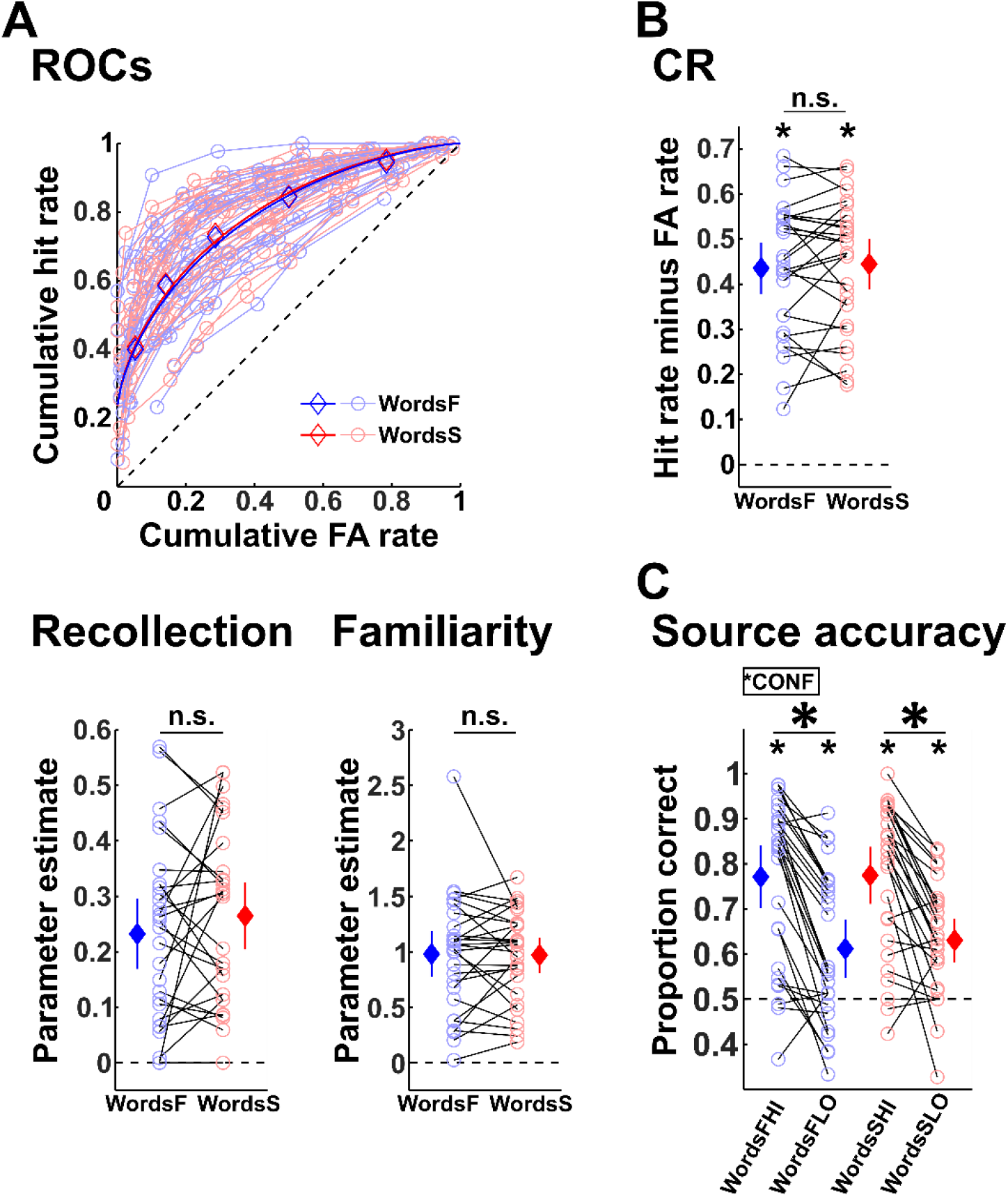
Behavioural results. Across memory measures, words previously associated with faces (wordsF) and scenes (wordsS) showed similar performance. **A**. Receiver operating characteristic (ROC) curves and behavioural modelling. The ROC plot (upper panel) depicts hit rates plotted against false alarm (FA) rates, cumulative over confidence levels. Note that all participants are above the chance diagonal. The fitted curves depict the DPSD predictions for wordsF and wordsS, here fitted to the group averages for illustrative purposes. The lower panel plots the single subject parameter estimates of the DPSD for recollection and familiarity. **B**. Corrected recognition (CR). **C**. Source accuracy. **Annotation:** Circles and line plots denote single participants; diamonds denote means across participants; error bars denote 95% confidence interval; dashed lines denote chance level. *CONF: significant main effect of recognition confidence. F: previous face association, S: previous scene association, HI: correctly recognised with high confidence, LO: correctly recognised with low confidence. \**p*<.05. n.s.: not significant.

First, we analysed participants’ recognition memory. During the scanned recognition phase, participants rated their recognition confidence for a given word on a scale of 1 (“sure new”) to 6 (“sure old”). From the distributions of hits and false alarms at each confidence level, we obtained receiver operating characteristic (ROC) curves, and estimated model parameters for recollection and familiarity (see Figure 1A). As this procedure assumes a lower bound of 0 for both parameters, we did not test them against zero (note that all single-subject ROC curves are above the chance diagonal). Recollection did not differ significantly between wordsF and wordsS (*t*_27_=1.124, *p*=.271), and neither did familiarity (*t*_27_=0.145, *p*=.885). Additionally, as a model-free measure of recognition performance, we computed corrected recognition (CR, hit rates minus false alarm rates, see Figure 1B). CR exceeded chance for both wordsF (*t*_27_=15.553, *p*<.001) and wordsS (*t*_27_=16.300, *p*<.001), and did not differ significantly between wordsF and wordsS (*t*_27_=0.578, *p*=.568).

In the post-scan source phase, for each word, participants gave a forced-choice response whether that word had been paired with a face or a scene the day before. Here, we analysed source accuracy for words that had been recognised with either high (HI) or low (LO) confidence during the recognition phase (i.e. HI: confidence rating = 6, LO: confidence rating = 4-5): wordsFHI, wordsFLO, wordsSHI, wordsSLO (see Figure 1C). A repeated measures ANOVA with the factors category and confidence revealed a highly significant main effect of confidence (*F*_(1,27)_=88.083, *p*<.001; no other effects, *p*≥.694) such that high-confident hits yielded higher subsequent source accuracy than low-confident hits. This confidence effect was confirmed using paired t-tests, with source accuracy greater for wordsFHI vs. wordsFLO, and for wordsSHI vs. wordsSLO (both *t*_27_≥5.652, *p*<.001). Finally, source accuracy exceeded chance for wordsFHI, wordsFLO, wordsSHI, and wordsSLO (all *t*_27_≥3.578, *p*≤.001).

### 2.2 fMRI results: Strategy

fMRI data were analysed within bilateral MTL subregion ROIs (HC, PRC, PHC, alEC, pmEC) that were manually delineated on the single-subject T1 scans (Insausti et al., 1998; Pruessner et al., 2000, 2002; Maass et al., 2015). First, we sought to establish category sensitivity during the perception phase, using both uni- and multivariate approaches. Then, only those ROIs showing such category sensitivity during perception, i.e. HC, PRC, and PHC (see Figure 2A), were considered for analyses of the recognition phase, as we were primarily interested in reinstatement of the perceptual activity. Here, we again employed both uni- and multivariate approaches.

**Figure 2.**
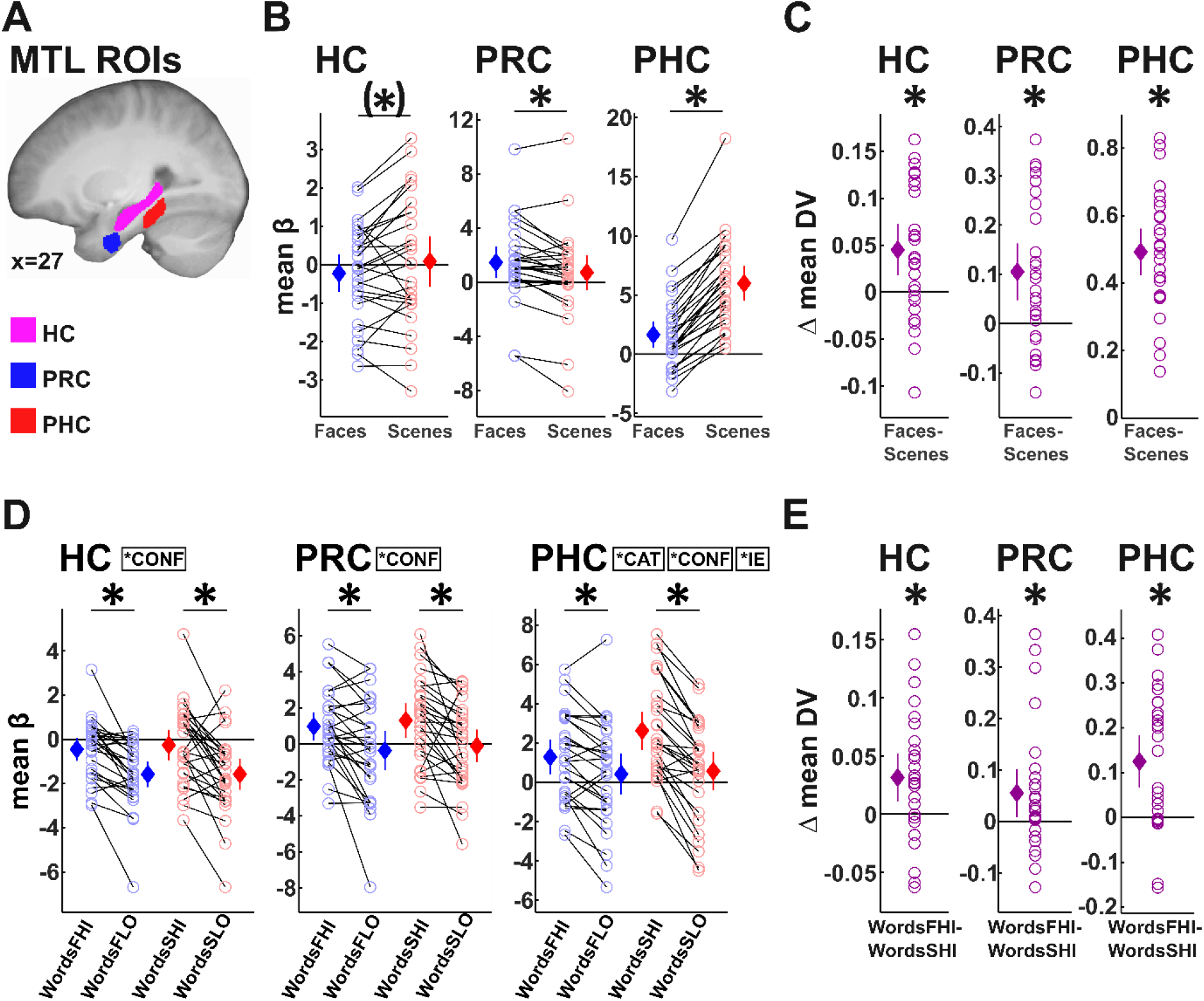
MTL subregions and fMRI results. **A**. For visualisation, single-participant ROIs of HC, PRC, and PHC were normalised, averaged across participants, thresholded at 0.5, and projected onto the mean normalised T1 image. **B**. Averaged beta values from the univariate analysis during face and scene perception. **C**. Differences between averaged decision values during face vs. scene perception from the multivariate decoding analysis. Positive difference values indicate discriminability of faces and scenes. **D**. Averaged beta values from the univariate analysis during high-vs. low confident correct recognition of words previously associated with faces vs. scenes. **E**. Results from the multivariate decoding analysis, with a classifier trained on face vs. scene perception, and tested on highly confidently recognised words previously associated with faces and scenes. Plotted are the differences between average decision values for wordsFHI and wordsSHI. Positive difference values indicate discriminability of wordsFHI and wordsSHI. **Annotation:** Circles and line plots denote single participants; diamonds denote means across participants, and error bars denote 95% confidence interval. *CAT: significant main effect of category; *CONF: significant main effect of recognition confidence; *IE: significant interaction effect of category and recognition confidence. F: previous face association, S: previous scene association, HI: correctly recognised with high confidence, LO: correctly recognised with low confidence. DV: decision value from the multivariate analysis. \**p*<.05, (*)*p*<.1.

### 2.3 Category sensitivity during perception

To establish category sensitivity during perception, we first analysed the MTL ROIs’ univariate response profiles by averaging beta estimates for the face and scene regressors from the perception phase within each ROI. A two-way repeated-measures ANOVA with the factors ROI (HC, PHC, PRC, alEC, pmEC) and category (faces, scenes) revealed a highly significant main effect of ROI (*F*_(2.80,75.57)_=16.341, *p*<.001) and category (*F*_(1,27)_=39.459, *p*<.001), as well as an interaction effect (*F*_(2.81,75.96)_=86.399, *p*<.001). Individual paired *t* tests within each ROI confirmed category sensitivity in PHC (scenes>faces, *t*_27_=13.593, *p*<.001) and PRC (faces>scenes, *t*_27_=3.400, *p*=.002), with a trend-level effect in HC (numerically scenes>faces, *t*_27_=1.878, *p*=.071) (see Figure 2B). There was no such effect in alEC or pmEC (both *t*_27_≤0.826, *p*≥.416). The PHC and PRC effects survived Holm-Bonferroni correction for multiple comparisons (5).

A multivariate decoding analysis complemented the univariate results. Multivariate analyses consider voxel patterns consisting of both activations and deactivations, thereby increasing sensitivity (Hebart and Baker, 2018). Face vs. scene classifiers were trained and tested on each ROI’s perception data in a leave-one-run-out fashion. We tested the differences between average decision values for face and scene trials (category discriminability) against zero. Category discriminability was above chance in HC, PRC, and PHC (HC: *t*_27_=3.379, *p*=.002, PRC: *t*_27_=3.739, *p*<.001, PHC: *t*_27_=14.595, *p*<.001, see Figure 2C), but not in alEC or pmEC (both *t*_27_≤0.389, *p*≥.700). The effects in HC, PRC, and PHC survived Holm-Bonferroni correction for multiple comparisons (5).

### 2.4 Category sensitivity during word recognition

Having established category sensitivity during perception in HC, PRC, and PHC, we turned to testing the word recognition data from these ROIs for effects of prior association with faces vs. scenes. As recollection-related neural reinstatement is thought to be restricted to high-confidence recognition (Yonelinas et al., 2010), we analysed the univariate response profiles of the MTL ROIs separately for high vs. low confidence hits, and for words previously associated with faces vs. scenes (wordsFHI, wordsFLO, wordsSHI, wordsSLO, see Figure 2D). A three-way repeated measures ANOVA with the factors ROI (HC, PRC, PHC), associated category (face, scene), and recognition confidence (high, low) revealed a significant three-way interaction of ROI, category, and confidence (*F*_(1.39,37.59)_=3.998, *p*=.040). Most other effects were also (marginally) significant (ROI: *F*_(1.74,46.92)_=13.684, *p*<.001; category: *F*_(1,27)_=6.501, *p*=.017; confidence: *F*_(1.27)_=59.648, *p*<.001; ROI x category: *F*_(1.66,44.93)_=5.240, *p*=.013; category x confidence: *F*_(1,27)_=4.165, *p*=.051; ROI x confidence: *F*_(1.52,41.05)_=0.292, *p*=.688). We followed up on the three-way interaction by computing separate two-way ANOVAs (category, confidence) within each ROI. All three ROIs showed highly significant main effects of confidence (all *F*_(1,27)_≥20.205, all *p*<.001). PHC additionally showed a main effect of category (*F*_(1,27)_=11.468, *p*=.002) and, critically, an interaction of category and confidence, with a larger confidence effect for words previously associated with scenes than with faces (*F*_(1,27)_=17.174, *p*<.001). There was no such interaction in HC or PRC (all *p*≥.525). Follow-up paired *t*-tests between wordsFHI vs. wordsFLO, and wordsSHI vs. wordsSLO were significant in all ROIs (all *t*_27_≥3.587, all *p*≤.0013), and all tests survived Holm-Bonferroni correction for multiple comparisons (6). In sum, all ROIs showed highly significant univariate activity increases during high-confident compared to low-confident correct word recognition, with the PHC particularly engaged during high-confident recognition of words previously associated with scenes.

Finally, we turned to our central analysis of multivariate decoding of the recognition data. The above univariate analysis is limited in that it focuses on overall activity differences between conditions, averaged across each ROI’s voxels. Multivariate analyses, in contrast, consider the information that is contained in each ROI’s activation pattern (Hebart and Baker, 2018). Here, because recollection is thought to involve reinstatement of associated information from the encoding context and lead to high confident hits (Eichenbaum et al., 2007; Yonelinas et al., 2010), we assume that neural activity during high-confident word recognition contains information about the previous face or scene association, reinstating patterns that were present during perception. Hence, a classifier trained to distinguish between faces and scenes during the perception phase and tested on high-confident hits during the recognition phase ought to be able to distinguish between previous face and scene associations. Thus, for each ROI, we tested category discriminability for high confidence words (i.e. the differences between average decision values for wordsFHI and wordsSHI) against zero (see Figure 2E). Category discriminability was above chance in all three ROIs (HC: *t*_27_=3.090, *p*=.005, PRC: *t*_27_=2.432, *p*=.022, PHC: *t*_27_=4.361, *p*<.001), and all three ROI effects survived Holm-Bonferroni correction for multiple comparisons (3).

## 3. Discussion

In the present study, we investigated whether high confident recognition of words is accompanied by incidental reinstatement of previously associated faces or scenes in subregions of the medial temporal lobe (MTL). During the recognition phase, participants rated their confidence that a given word was old or new, but, critically, were not asked to intentionally retrieve associated categorical information. Behaviourally, participants successfully recognised words from the encoding phase, and their subsequent source memory for associated categorical information was above chance. Analysis of the fMRI data first confirmed category sensitivity during perception in the MTL. Second, importantly, our data revealed incidental category reinstatement in MTL subregions during word recognition: Hippocampus (HC), perirhinal cortex (PRC), and parahippocampal cortex (PHC) demonstrated multivariate discriminability of previous faces vs. scene associations, using a classifier trained on the perception data. Crucially, the perception and recognition phases did not share any perceptual input, as the perception phase presented faces and scenes, but not words, and the recognition phase presented words, but not faces or scenes. Hence, these multivariate results can only reflect reinstatement of associated face and scene information during word recognition. In addition, the univariate activity profiles of the MTL ROIs during the recognition phase showed robust activity increases for high compared to low confident words.

The present study provides novel evidence for incidental reinstatement of faces and scenes in the MTL in a word recognition task. This is in line with the dual process signal detection model (DPSD) of recognition memory, which assumes that some of the queried words – namely, those that are recollected – are accompanied by neural reinstatement of the associated information (Eichenbaum et al., 2007; Yonelinas et al., 2010). Indeed, our data demonstrate that words recognised with high confidence show such reinstatement in the MTL by allowing for multivariate decoding of the previously associated category. This observed category sensitivity within the MTL follows from its anatomical connectivity patterns. To simplify, the PRC vs. PHC serve as relay stations for object-related vs. spatial information, respectively, between the visual system and the HC (Davachi, 2006; Eichenbaum et al., 2007). This account is exemplified in our univariate perception results, with enhanced activity during face perception in the PRC, and enhanced activity during scene perception in the PHC. A number of previous fMRI studies have demonstrated such category dissociations between PRC and PHC during perception and encoding of faces (or objects) vs. spatial stimuli (or scenes) (Awipi and Davachi, 2008; Litman et al., 2009; Staresina et al., 2011; Liang et al., 2013; Berron et al., 2018; Schultz et al., 2019, 2021). Importantly, the bidirectionality of the underlying MTL connectivity might support the reinstatement of representations during memory retrieval (Davachi, 2006; Eichenbaum et al., 2007). Indeed, MTL content sensitivity in the absence of perceptual input, implying cortical reinstatement, has been demonstrated previously (Schultz et al., 2012, 2019; Staresina et al., 2013; Mack and Preston, 2016; Liang and Preston, 2017). Note that these studies investigated intentional retrieval – e.g. Schultz et al. (2019) presented words and asked participants to vividly retrieve a previously associated object vs scene, which was associated with i) elevated category-sensitive univariate retrieval activity in PRC vs. PHC, and ii) increased across-voxel correlation of category-sensitive retrieval and perceptual activity.

In contrast, and complementing these earlier results, we investigated reinstatement during recognition. Participants rated their recognition confidence for a given target or distractor word, but were not instructed to retrieve the associated category information. To our knowledge, no previous study has investigated cortical reinstatement in MTL subregions in a recognition memory paradigm without explicit instruction to retrieve associated information. Two studies (Skinner et al., 2014; Bowen and Kensinger, 2017) had participants give recognition judgments for words previously paired with faces and scenes without explicit instruction to retrieve the associated information; however these studies did not focus on subregions of the MTL. Another study (Kuhl et al., 2013) also presented words that had been previously learned with faces and scenes. However, this was not a word recognition task: Participants were asked to explicitly retrieve information about the associated images, either the category of the image (face or scene), or its location (left or right). Here, MTL retrieval representations tracked category regardless of whether participants were focussing on the category or location of the image they were retrieving (however, their ROIs did not distinguish between PRC, PHC, and EC). Note that the absence of an explicit instruction to retrieve associated information in our study does not imply that recollection of these associations was non-conscious, or that the participants actively suppressed these associations. Furthermore, the subsequent behavioural test of source memory indicates that they had retained above-chance explicit memory for the associated category information. Our results demonstrate that cortical reinstatement in the MTL does not require an instruction of intentional, vivid retrieval.

Whereas our multivariate results give clear evidence for category reinstatement in the MTL, the univariate data are dominated by category-insensitive confidence effects across MTL subregions. Only in PHC were these effects increased for one category (scenes). The uni- and multivariate approaches differ on a number of dimensions. First, multivariate analyses are generally thought of as more sensitive than univariate analyses (Hebart and Baker, 2018), which may explain why the multivariate analysis yielded evidence for category reinstatement in PRC while the univariate analysis did not. Moreover, while univariate analyses assume that a ROI’s involvement in a process will be reflected in elevated mean activity, multivariate analyses assume that both activations and deactivations equally contribute to the information represented in a given ROI (Hebart and Baker, 2018). Here, some caution is warranted regarding the interpretation of our multivariate results: Given the univariate activity differences during the perception phase, a parsimonious explanation would be that, during recognition, PRC represents the retrieved face information, while PHC (and HC) represent the retrieved scene information. This would be in line with earlier functional reports (Schultz et al., 2012, 2019; Staresina et al., 2013; Mack and Preston, 2016) as well as the PRC’s and PHC’s anatomical connectivity to regions of the ventral and dorsal visual stream, respectively (Suzuki and Amaral, 1994a; Eichenbaum et al., 2007). However, in our multivariate analyses, evidence for faces cannot be distinguished from evidence against scenes, and vice versa. This means that each ROI’s ability to discriminate between face and scene associations may be driven by that ROI representing face information, or scene information, or both. Hence, our results imply that PRC, PHC, and HC maintain information about previous associations of the words during the recognition phase, but based on the multivariate results alone, we cannot conclude a preference of one category over the other. Indeed, as scenes typically contain objects, to which PRC is sensitive (Robin et al., 2019), it is likely that scene reinstatement also engages PRC to some degree. Finally, as the multivariate analysis classifies recognition trials based on neural patterns from the perception data, it is a direct test of the reinstatement concept, which implies topographical and informational overlap between perception/encoding and retrieval (Danker and Anderson, 2010). Recent approaches, however, have also investigated differences between encoding- and retrieval-related memory representations, emphasising shifts in representational granularity and brain topography (Baldassano et al., 2016; Bainbridge et al., 2020), direction of the information flow (Staresina et al., 2013; Linde-Domingo et al., 2019), and transformation of the memory trace itself (Xiao et al., 2017; Favila et al., 2018). Reinstatement, as investigated here, is therefore only one facet of how the brain represents the content of memory during retrieval.

In the univariate data, we observed category-sensitive effects of recognition confidence in PHC, but, unexpectedly, not PRC. This is in contrast to earlier studies showing category-sensitive univariate effects in PRC and PHC during intentional, vivid retrieval (Schultz et al., 2012, 2019; Staresina et al., 2013). Given that all our behavioural measures indicate comparable memory performance for words previously associated with faces and scenes, this effect cannot be attributed to memory performance differences across conditions. However, recent results suggest that scenes could be special memoranda compared to e.g. faces or objects, increasing associative memory by providing a spatial context that binds to items more easily than other material (Robin and Olsen, 2019). Furthermore, although eliciting comparable memory performance, the scenes in our stimulus set had more diverse content (e.g. a mountain side, a coast, a forest) than the face stimuli. These properties could have increased scene reinstatement during the word recognition task. It is important to point out that PRC does not only receive information from the ventral visual stream, but additionally receives a number of inputs from spatial processing regions such as the PHC (Suzuki and Amaral, 1994a; Burwell and Amaral, 1998b). Accordingly, studies have reported evidence for similar processing of object-related and spatial information in the PRC under some circumstances (Berron et al., 2018; Lawrence et al., 2020).

Our results show category discriminability in the HC for both the perception and recognition phases, as well as (marginally) elevated mean activity during viewing of scenes compared to faces. Some previous studies, including both functional imaging and lesion studies, imply scene specialisation in the HC (Lee et al., 2005a, 2005b; Graham et al., 2006; Taylor et al., 2007; Zeidman et al., 2015), in line with a prominent role of the HC in spatial processing (Maguire and Mullally, 2013; Maguire et al., 2016; Bellmund et al., 2018). Other studies, however, have shown no evidence for category-level distinctions in the HC (Staresina et al., 2012, 2013; Mack and Preston, 2016; Schultz et al., 2019). Anatomy-based models of MTL function imply a primarily associative or relational role of the HC in episodic memory; in this view, the HC is insensitive to stimulus category (Davachi, 2006; Eichenbaum et al., 2007). However, commonalities between relational, or associative, and spatial hippocampal processing have been noted (Buzsáki and Moser, 2013; Eichenbaum, 2017). Future work may illuminate the circumstances under which HC scene preferences prevail.

We note some limitations of the current study. First, recent years have seen rising interest in the role of anterolateral and posteriormedial EC subregions (alEC, pmEC) during perception/encoding and retrieval, establishing the notion of category sensitivity in EC subregions during these processes (Schultz et al., 2012, 2015; Reagh and Yassa, 2014; Maass et al., 2015; Navarro Schröder et al., 2015; Berron et al., 2018). Here, we found no evidence for category-sensitive representations in the EC. One methodological challenge in fMRI of the MTL is the signal quality gradient from anterior to posterior MTL, leading to decreased signal-to-noise ratio and increased susceptibility artifacts in the anterior MTL cortex, including the EC (Carr et al., 2010). Hence, our null-results in the EC may be due to signal quality issues. A second potential limitation concerns the recollection vs. familiarity distinction. According to the DPSD, high-confident hits may consist of both recollected and highly familiar items (Yonelinas et al., 2010). While we assume that, based on the underlying model, the observed sensitivity to the associated category during recognition memory was driven by the recollected items rather than the highly familiar items (Eichenbaum et al., 2007; Yonelinas et al., 2010), these processes cannot be disentangled on the item level. Hence, we cannot rule out incidental reinstatement for highly familiar items. Lastly, there has been a discussion whether the MTL is involved in perception at all, or whether all MTL processing is necessarily mnemonic (Suzuki and Baxter, 2009). Our results do not resolve this debate. While we have treated the category-specific MTL processes during the perception phase as perceptual in nature, we cannot rule out that they have a primarily mnemonic function, i.e. memory encoding (Awipi and Davachi, 2008; Staresina et al., 2011).

In summary, we show that, in the absence of any differences in perceptual input, the three major subregions of the MTL – HC, PRC, and PHC – nonetheless contain representations of associated category information (faces/scenes) during word recognition. These data support a functional model of episodic memory in the MTL that is informed by anatomical connectivity, and that emphasises the similarity of content representations during perception/encoding and retrieval. Future work may clarify the role of human entorhinal subregions during long-term retrieval of category-sensitive representations, as well as differences in representations involved during perception/encoding vs. retrieval.

## 4. Materials and Methods

### 4.1 Sample

We report data from 28 volunteers (8 male, mean age 26.0, range 18-35). Three more were excluded from data analysis (one for excessive head movement, one for falling asleep in the scanner; one dropped out after day one). All volunteers were right-handed, healthy with normal or corrected-to-normal vision, and reported no past neurological or psychiatric diagnoses. They received monetary reimbursement of €10/hour + bonus. Prior to participation, they gave written informed consent. The study procedure was approved by the local ethics committee (Hamburg Board of Physicians).

### 4.2 General procedure

The experiment was conducted over two consecutive days. Day one comprised the scanned perception (∼30min) and unscanned encoding phase (∼40min). Day two comprised the scanned recognition (∼59min), and unscanned source phase (∼25-40min). Mean lag between perception and recognition phase was 19.5h (range: 13.5-24h).

### 4.3 Stimuli

Stimuli consisted of 10 grayscale neutral faces (Endl et al., 1998) and 10 grayscale natural outdoor scenes (various internet sources) used in a previous study (Schultz et al., 2012), as well as 390 emotionally neutral, concrete German nouns from the Berlin Affective Word List Reloaded (Võ et al., 2009). For each participant, 260 words were randomly selected as encoding items, whereas the remaining 130 served as distractors during the recognition phase.

### 4.4 Behavioural tasks

The *perception phase* was a modified Monetary Incentive Delay (MID) task (Knutson et al., 2000) (see Figure 4A for details). Each of the 20 faces and scenes appeared 6 times, resulting in 120 trials over 3 runs. Trial order was pseudo-randomised, with no more than 4 face or scene trials appearing in a row. Trials started with a 1€ coin followed immediately by a face or scene (initial image presentation). At a random timepoint during image presentation, a flicker (blank screen for 1 frame) prompted participants to press a button as fast as possible to win a reward, using a button box held in their right hand. Finally, the image reappeared with reward feedback. Response time limits adapted to a reward probability of 0.735 over trials, separately for faces and scenes. Participants received a bonus for each earned reward, amounting to ∼€3.20 in total.

**Figure 4.**
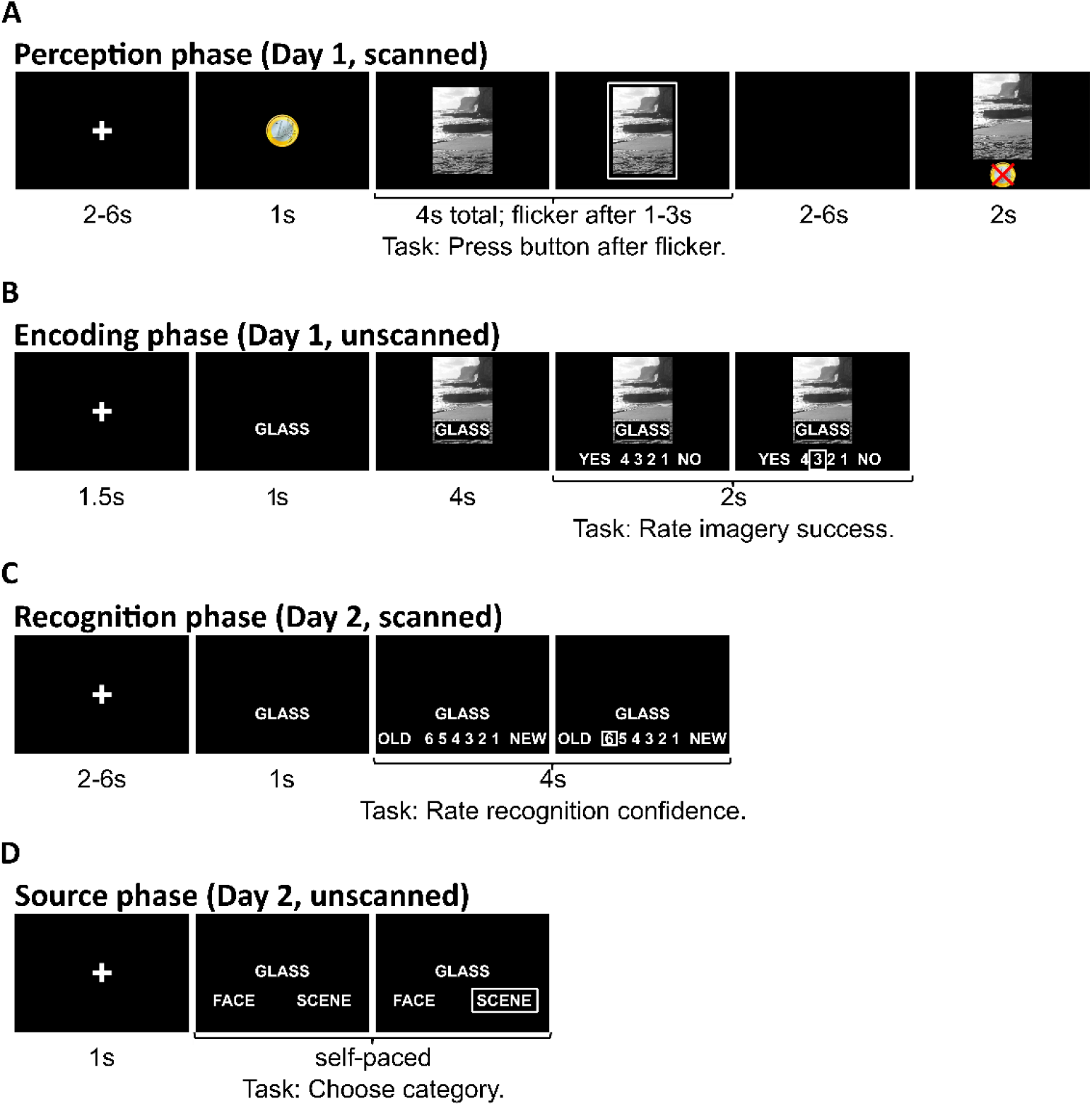
Example trials for the experimental phases. A: Perception phase. B: Encoding phase. C: Recognition phase. D: Source phase.

In the *encoding phase* (see Figure 4B for details), participants encoded associations between words and images. Each of the 20 faces and scenes was paired with 13 words, resulting in 260 trials, presented in 5 blocks with self-paced breaks. Trial order was pseudorandomised, with no more than 4 face or scene trials in a row, and identical images separated by at least 10 trials. Per trial, participants were asked to combine a word and image into a single mental image, and rated their imagery success on a scale of 1 to 4. Response layout (1-2-3-4 or 4-3-2-1) was randomly switched between trials.

In the *recognition phase* (see Figure 4C for details), all 260 encoded words plus 130 distractor words were presented over 5 runs. Trial order was pseudorandomised so that no more than 4 distractors and 4 words associated with either a face or a scene appeared in a row. Per trial, participants were asked to rate their confidence that a given word was new or old, on a scale of 1 to 6. Participants used two button boxes held in their left and right hand, with three buttons mapped on each. Response layout (1-2-3-4-5-6 or 6-5-4-3-2-1) was randomly switched between trials. No instruction was given to retrieve the associated image.

In the *source phase* (see Figure 4D for details), all 390 words were presented again, and participants indicated whether a given word had been paired with a face or a scene the day before. Additionally, they indicated whether they associated the word with a reward or not (not pictured; data not included in the present report).

All tasks were programmed using Presentation® software (Version 14.9, Neurobehavioral Systems, Inc., Berkeley, CA, www.neurobs.com).

### 4.5 Behavioural analyses

For the recognition phase, we analysed a model-free outcome measure (corrected recognition, CR) as well as two model-based outcome measures (recollection, familiarity). CR was computed as the difference between hit rate minus false alarm rate (the proportions of confidence ratings ≥ 4 for targets minus distractors, respectively). Recollection and familiarity are parameters in the dual-process model. This model assumes that two processes contribute to recognition memory: An all-or-none threshold process (recollection), and a signal detection process (familiarity) (Yonelinas et al., 2010). Parameter estimates for these processes were obtained from each participant’s distribution of recognition confidence ratings, separately for wordsF and wordsS, using maximum likelihood estimation (Dunn, 2010). Finally, for the source phase, we computed source accuracy (proportion of correctly identified source category) for words that, during the recognition phase, had been correctly recognised with high confidence (confidence rating = 6) respectively low confidence (confidence rating = 4-5), separately for wordsF and wordsS.

### 4.6 MR data acquisition

MR data was acquired on a 3T Siemens TIM TRIO scanner using a 32 channel head coil. The perception and recognition phases were scanned using a high-resolution T2*-weighted EPI sequence (33 descending slices, no gap, 1.5mm isotropic voxels, TR=2.49s, TE=30ms, PAT factor 2) with the field of view aligned to the longitudinal MTL axis. On day 2, an additional T1-weighted MPRAGE structural scan was acquired (240 slices, 1×1×1mm).

### 4.7 ROI approach

All statistical analyses were conducted in single-subject space within bilateral masks of MTL subregions (HC, PRC, PHC, alEC, pmEC) that were manually segmented on the T1 following existing guidelines (Insausti et al., 1998; Pruessner et al., 2000, 2002; Maass et al., 2015) using MRIcron (Rorden and Brett, 2000). For PRC and PHC, the middle third of the parahippocampal gyrus (i.e. posterior PRC and anterior PHC) was discarded to maximise category sensitivity within these regions (Staresina et al., 2013; Schultz et al., 2019).

### 4.8 MR data preprocessing

MRI data were analysed in Matlab/SPM12 except where noted. Functional images were corrected for slice acquisition time, head movement, and movement-related distortions. T1 images were coregistered to the functional data using boundary-based registration (FSL epi_reg). In order to create Figure 2A, T1 images were segmented into grey matter, white matter, and cerebrospinal fluid, and the resulting flowfields were used to normalise the T1 images and single-subject ROIs into MNI (Montreal Neurological Institute) space.

### 4.9 Univariate analyses

For the univariate analyses, we set up two categorical first-level linear models (GLMs) on the non-normalised, unsmoothed data from the perception and recognition phase, respectively. Runs were concatenated within each phase. Regressors were modelled as stick functions convolved with the canonical hemodynamic response function. For the perception phase, regressors of interest comprised faces vs. scenes during initial image onset, when no reward information was available. Also modelled were the reward feedback onsets, separately for face/reward, face/no reward, scene/reward, and scene/no reward. For the recognition phase, regressors were modelled on the word onset, and regressors of interest comprised words previously associated with faces (F) vs. scenes (S) and correctly recognised with high (HI, confidence rating = 6) vs. low confidence (LO, confidence rating = 4-5) (wordsFHI, wordsFLO, wordsSHI, wordsSLO). Also modelled were misses, separately for wordsF and wordsS (confidence ratings ≤ 3), false alarms and correct rejections for the distractor items, and error trials. Models included a high-pass filter (128s) and autoregressive model (AR(1)) as well as run constants. Beta values from the regressors of interest were averaged within each ROI and submitted to group-level analyses.

### 4.10 Multivariate analyses

The first-level GLMs underlying the multivariate analyses were set up similarly to the univariate analyses, albeit with a single-trial regressor on each initial image onset as well as reward feedback onset (perception phase, the latter were discarded), and on each word onset (recognition phase). Multivariate decoding was applied to single-trial *t*-values from each ROI (z-scored within each voxel separately for training and test data), using regularised linear discriminant analysis (LDA) as implemented in the MVPA-light toolbox (Treder, 2020). Two decoding analyses were conducted: First, we tested category discriminability during perception. Here, we trained and tested a classifier on face vs. scene perception in a leave-one-run-out fashion. Second, we tested category reinstatement during the recognition phase. Here, we trained a classifier on face vs. scene perception during the perception phase, and tested it on high-confident hits for words previously associated with faces vs. scenes (wordsFHI vs. wordsSHI) from the recognition phase. These analyses resulted in decision values for each testing trial and ROI, with increasing values indicating face evidence, and decreasing values indicating scene evidence. As the zero point in these analyses is arbitrary (representing the midpoint of all trials in the testing set), we computed group level analyses on the differences between decision values for face minus scene trials (perception phase) and wordsFHI minus wordsSHI (recognition phase), with positive difference values indicating face vs. scene discriminability.

## 5. Acknowledgments

We thank Matthias Treder for helpful discussions of the multivariate analysis approach.

## Footnotes

1 Note that this experiment was originally devised to additionally assess the influence of associated reward on cortical reinstatement. Hence, the perception phase was designed as a reward task. As reward did not have reliable effects on behavioural or neural measures of memory, we here reanalyse the dataset omitting this factor. The brain responses extracted from this phase were modelled at a timepoint in each trial in which no reward information was available.

